# *β*-cells operate collectively to help maintain glucose homeostasis

**DOI:** 10.1101/765933

**Authors:** Boris Podobnik, Dean Korošak, Maša Skelin Klemen, Andraž Stožer, Jurij Dolenšek, Marjan Slak Rupnik, Plamen Ch. Ivanov, Petter Holme, Marko Jusup

## Abstract

Residing in the islets of Langerhans in the pancreas, beta cells contribute to glucose homeostasis by managing the body’s insulin supply. A circulating hypothesis has been that healthy beta cells heavily engage in cell-to-cell communication to perform their homeostatic function. We provide strong evidence in favor of this hypothesis in the form of (i) a dynamical network model that faithfully mimics fast calcium oscillations in response to above-threshold glucose stimulation and (ii) empirical data analysis that reveals a qualitative shift in the cross-correlation structure of measured signals below and above the threshold glucose concentration. Combined together, these results point to a glucose-induced transition in beta-cell activity thanks to increasing coordination through gap-junctional signaling and paracrine interactions. The model further suggests how the conservation of entire cell-cell conductance, observed in coupled but not uncoupled beta cells, emerges as a collective phenomenon. An overall implication is that improving the ability to monitor beta-cell signaling should offer means to better understand the pathogenesis of diabetes mellitus.

---

On a fundamental level, living tissues comprise genetically identical cells of the same differentiation fate that work cooperatively to support homeostasis and other physiological functions [1]. Cooperation, however, is susceptible to cheating unless there is a mechanism for mutual recognition of cooperators referred to as positive assortment [1]. Cell-to-cell communication is perhaps the most obvious means of positive assortment, employed even by cancer cells—quintessential cheaters from the perspective of normal tissue functioning—in order to successfully metastasize [2]. A key question in this context is how, in large clusters of cells (e.g., tissues or organs), cluster-wide functionalities emerge as collective phenomena from local cellular interactions (e.g., communication and cooperation) encoded into individual cells. Endocrine cells in pancreatic islets, in particular, form well-defined cellulo-social clusters, each about 100 *μ*m in size. Intriguingly, islet size is persistent across multiple vertebrate species [3], thus fueling conjunctures that cross-species size persistence plays a vital role in the collective functioning of a healthy islet [4].

Beta cells are a type of pancreatic cells that participate in glucose homeostasis by producing, storing, and releasing the hormone insulin [5–7]. When the glucose blood level is high, a state known as hyperglycemia, beta cells respond with a release of insulin into the bloodstream to promote nutrient transport into a variety of cells in support of anabolic metabolism. When the glucose blood level is low, a state known as hypoglycemia, or glucose is otherwise in demand, e.g., due to physical activity, beta cells switch off insulin secretion to allow catabolic metabolism. Beta cells thus employ a negative feedback mechanism to help maintain the blood glucose within a safe range. In comparison, therapeutic administration of insulin in diabetic patients lacks the precision of the internal regulatory processes, yielding either too high or too low plasma glucose levels [8]. Hypoglycemic episodes arising in relation to insulin therapy are a common adverse effect of antidiabetic therapies [9]. Although the basic outline of glucose homeostasis is conceptually clear, complexity quickly escalates when looking for a microscopic understanding of the negative feedback mechanism employed by beta cells. We herein attempt to better understand this complexity, focusing in particular on cell-to-cell communication in the face of mounting evidence that such communication is important for insulin secretion and, by extension, glucose homeostasis [6–8].

Illustrating microscopic complexity, cellular nutrient-sensing mechanisms [10, 11] have been shown to intermix metabolic signals, electrical activity, and cytosolic calcium signaling [12, 13]. This complex intermix prevents beta cells of a healthy pancreas to oversecrete insulin despite huge intracellular insulin stores [14], which exceed the lethal dose two to three orders of magnitude if released at once. A possible answer to how beta cells regulate their secretion may lie in a response of betacell collectives to above-threshold glucose concentrations (>7 mM in mice [15, 16]) at which electrical and calcium activities flip between non-stimulatory and stimulatory phases. Here, the term collective is used to denote the fact that beta cells, in addition to engaging in paracrine interactions [17, 18], are coupled to neighboring beta cells via gap junctions (comprising two connexons of Cx36 protein) to form a communicating functional syncytium [19]. This functional syncytium as a whole reacts to nutrients (e.g., glucose) or pharmacological substances (e.g., sulphonylureas) in a fundamentally different manner from isolated cells [20] or partially ‘uncoupled’ cells, i.e., those lacking the Cx36 protein [21]. Specifically, after closing ATP-sensitive K^+^ channels in a coupled configuration, the remaining entire cell-cell conductance varies among individual cells, but stays constant and relatively high islet-wide. This happens throughout the time-frame of a typical experiment (>10 minutes) and despite the concurrent dynamic interchange of non-stimulatory and stimulatory collective phases [21].

A series of models describing fast Ca^2+^ oscillations in the electrical activity of beta cells have been constructed since the first minimal model over three decades ago [22]. The primary purpose of these models, however, has been to generate the saw-tooth time profile characteristic of slow processes underlying the observed electrical bursting [23, 24]. By contrast, there seem to have been much less concern with the oscillatory response of beta cells to the threshold glucose concentration, and the relationship of this response to communication within the functional syncytium or the conservation of entire cell-cell conductance.

We demonstrate that a cell-to-cell communication model based on a dynamical network approach [25, 26] captures the essential features of fast Ca^2+^ signaling, and that within such a model the simulated equivalent of the entire cell-cell conductance remains conserved in a statistical sense after the network transitions from an inactive to an active state, thus suggesting that the same mechanism may be at work in a living cell system as well. To provide empirical support for these results, we proceed with probing correlations in fast Ca^2+^ signaling as a measure of cell-to-cell communication, and present evidence of a shift in the correlation structure between non-stimulatory and stimulatory conditions. This is consistent with the hypothesis that the living cell system, just as the modeled dynamical network, attains sensitivity to threshold glucose stimulation via collective action.

## DATA CHARACTERIZATION

We analyzed a dataset obtained by Ca^2+^ imaging of acute pancreatic tissue slice [27] comprising rodent pancreatic oval-shaped islet (approx. dimensions: 370μm×200μm). We recorded Ca^2+^ signals with a functional multi-cellular imaging technique at 10 Hz and 256×256 pixel resolution in 8-bit grayscale color depth. The dataset consisted of 65,536 traces of calcium signals, each 23,873 time steps long. During the recording, we increased glucose concentration in the chamber containing the tissue sample from 6 mM to 8 mM and then decreased back to initial concentration near the end of the experiment. We chose these two physiological glucose concentrations to induce a transition from a non-stimulatory to a stimulatory beta-cell phase. The typical threshold for this transition in mice is around 7 mM [15, 16]. Further methodological details, including an Ethics Statement, are available in Supplementary Methods.

Individual Ca^2+^ traces, as well as the ensemble-averaged signal, exhibit (i) slow calcium oscillations with a period of approximately 5 min, but also superimposed (ii) fast calcium oscillations (Fig. 1A). We detrended all traces as a part of signal preprocessing to exclude the effect of systematic slow diminishing of the signal with time (Supplementary Fig. 1). Calcium oscillations on both timescales are known to be accompanied with insulin release [28], but the saturation kinetics of the Ca^2+^-dependent insulin release suggests that faster oscillations dominate [29]. The relation between slow calcium oscillations and pulsating insulin release was comprehensively described in a recent review [24]. Here, by contrast, we focus on fast calcium oscillations, which in acute pancreatic slice preparations, like in early micro-electrode recordings [30], directly correspond to membrane electrical bursting activity [16] and are thought to be functionally relevant for glucose homeostasis [31].

**Figure 1.**
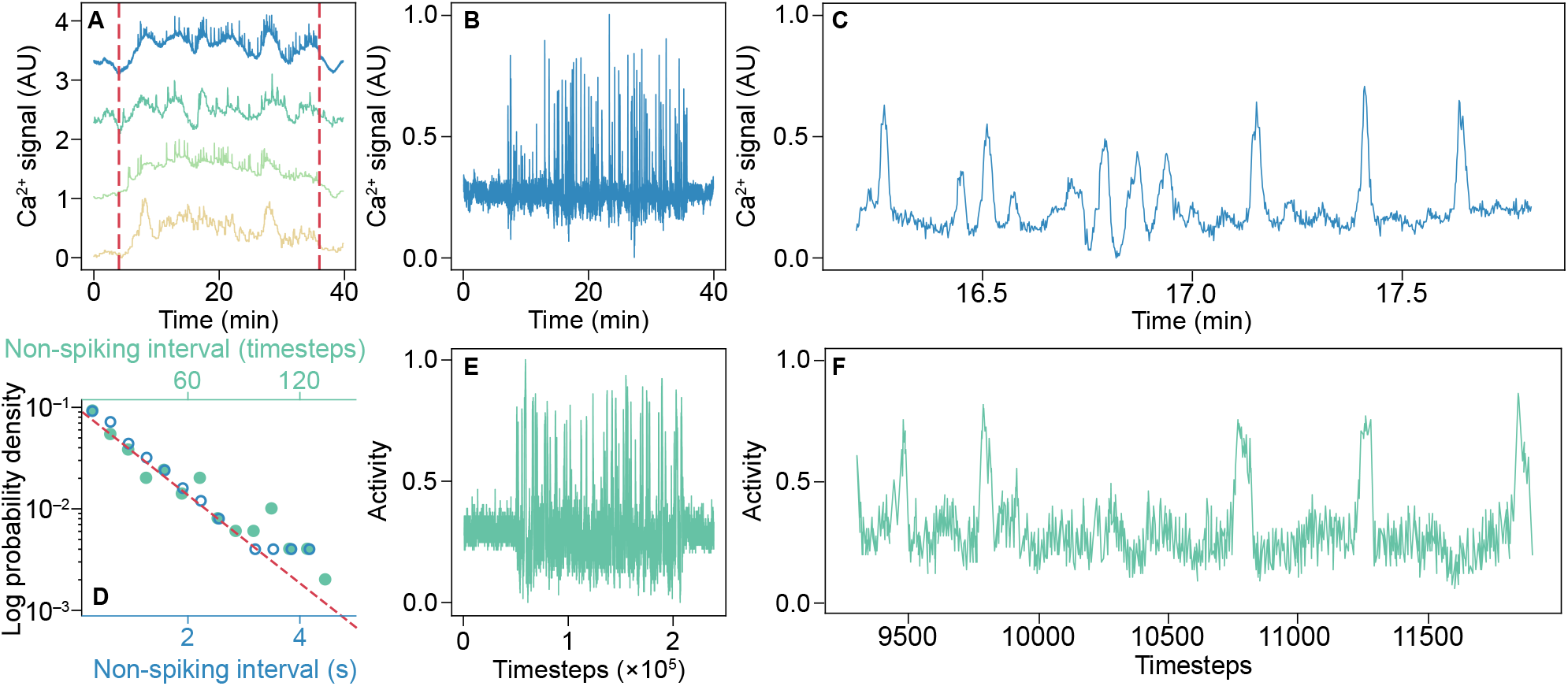
Beta-cell Ca^2+^ signaling is faithfully mimicked by a cell-to-cell communication model. **A**, Examples of the detrended experimental traces of Ca^2+^ signaling, including the ensemble average (top trace). Vertical red lines denote the time interval during which we perfused the tissue slice with 8 mM glucose solution. Otherwise, we kept the tissue slice in 6 mM glucose solution, which is below the typical threshold for glucose-stimulated activity of beta cells. **B**, The ensemble average of the measured fast Ca^2+^ signaling. The transitions from a non-stimulatory glucose to a stimulatory glucose phase and vice versa are clearly visible. **C**, A detail of the ensemble average of the measured fast Ca^2+^ signaling in the stimulatory phase. Spikes of exceptionally high activity are interspersed among somewhat quieter periods. **D**, The probability distribution of non-spiking intervals. Open dots represent the empirical non-spiking distribution for the ensemble-averaged fast Ca^2+^ signaling component in a semi-log plot. The dashed line is a theoretical distribution function, *ρ* exp(−*ρt*), with mean spiking rate *ρ* = 1 s. Among our primary goals was to reproduce the described empirical patterns using a cell-to-cell communication model (details in the main text). Solid dots thus represent the non-spiking distribution for the simulated fast beta-cell activity. **E**, Our cell-cell communication model with N = 100 cells successfully mimics fast Ca^2+^ signaling (cf. panel **B**). At transitioning from a non-stimulatory (6 mM) to a stimulatory (8 mM) glucose solution in the experiment, we set the model’s forcing parameter, i.e., cell responsivity, to a higher value. **F**, A detail of the simulated fast beta-cell activity. Despite a somewhat higher baseline noise compared to measurements in this particular example (cf. panel **C**), the two signals are statistically equivalent (see also Supplementary Fig. 3).

To isolate fast calcium oscillations, we proceeded by removing slow oscillations from traces using a total variation based filter [32]^1^ (Supplementary Fig. 2). The structure of the remaining fast calcium signal (Fig. 1B) clearly separates the non-stimulatory phase with little calcium activity from the stimulatory phase with large oscillation amplitudes. In both phases, the signal is stochastic in nature, which can be analyzed using statistical measures. We thus binarized the fast component of the ensemble-averaged Ca^2+^ trace (spiking or not; Fig. 1C), and subsequently calculated the non-spiking intervals. We found that the non-spiking intervals are governed by a Poisson process, implying that the probability density function (pdf) of the non-spiking intervals can be approximated with exponential distribution *ρ* exp(−*ρt*), where the mean spiking rate equals *ρ* = 1 s (open dots in Fig. 1D). Interestingly, a dynamical network model of cell-to-cell communication produces a simulated fast signal with the same statistical characteristics (solid dots in Fig. 1D). We next describe this model and some of its basic properties.

## EMERGING PROPERTIES OF BETA CELL-TO-CELL COMMUNICATION

Glucose homeostasis is a complex phenomenon involving multiple cell types and hormones. In our stylized model, however, we focus on beta cells alone, and in particular the role of cell-to-cell communication in shaping the response of these cells to the threshold glucose concentration. To this end, we started from a general dynamical network model [25] that explains how mutually interacting network elements conspire to cause highly non-linear dynamics with spontaneous phase flipping. We adapted the model in such a way that *N* network nodes represent individual beta cells and network links represent couplings with *k* neighboring cells. Each node can be in one of the two states, inactive or active, reflected in the amplitude of fast calcium signal. The state of the network in a particular time step is characterized by the fraction of active nodes.

The model’s key assumption is that nodes change their state internally or externally, where the former is modulated by the presence or absence of stimulatory glucose concentration, while the latter is due to cell-to-cell communication. An internal transition of a node from active to inactive state over time period *dt* occurs with probability *pdt*. In contrast, a transition prompted by mutual communication occurs with probability rdt, but only if the node has less than *m* < *k* active neighbors. Nodes return to their original state after relaxation times *τ_i_* and *τ_e_* following an internal or an external transition, respectively.

Once node relaxation is defined, the network’s state is determined by two parameters, probability rate r and the average fraction of internally inactive nodes, which follows from probability rate *p* via *p*\* = 1 – exp(-*pτ_i_*) [25]. In two-parameter phase space, (*r, p*\*), there is a critical point that opens a bi-stability region separating low- and high-activity model equilibria [25, 26]. When network parameters approach the critical point, node activity suddenly picks up, whereas inside the bi-stability region even spontaneous phase flipping becomes possible [25, 26].

To illustrate similarities between model outputs and recorded Ca^2+^ traces (cf. panels B, C and E, F in Fig. 1), we ran exhaustive numerical simulations (Supplementary Remark 1). First, we created a random regular network with *N* = 100 nodes, each with *k* = 10 neighbors, where the threshold for external inactivation was set to *m* = 4. We then chose *r* = 0.78, *τ_i_* = 10, and *τ_e_* = 1 as the parameter values that faithfully reproduce the characteristics of recorded Ca^2+^ traces (Supplementary Fig. 3). Finally, we began simulations with *p*\* = 0.90 to place the network into the inactive part of the phase space, but subsequently decreased this value to *p*\* = 0.28 to position the network close enough to the critical point for the activity to pick up substantially (Supplementary Fig. 4; see also Supplementary Fig. 5 for some additional model properties near the critical point). The decrease of *p*\* follows a decrease in probability rate *p*, which in turn is a response to an increased glucose concentration. As mentioned, the presence or absence of stimulatory glucose concentration modulates the response of individual beta cells, while the response of the beta-cell collective remains muted until a certain critical threshold of individual cell activity is reached. This behavior mimics the idea, first proposed by modeling studies [33] and then empirically validated [21], that the concentration response of an average electrical activity in beta-cell collectives is steep with respect to glucose sensing. Cell-cell coupling thus narrows the glucose concentration range that induces or stops insulin secretion, which is in sharp contrast with Cx36-deficient mice whose inability to synchronize beta-cell activation, activity, and deactivation leads to increased basal insulin release [21, 34].

As with the recorded fast calcium oscillations, we calculated the pdf of non-spiking intervals for the binarized network activity and found the same Poisson process as in the recorded data (cf. open and solid dots in Fig. 1D). This calculation, in particular, shows that the similarities between measurements and simulations are not just superficial, but extend into the statistical domain as well.

To strengthen the case for similarity between measurements and simulations in the statistical domain, we compared additional statistics implied by the data and predicted by the model. To this end, we first differenced both measurements and simulations to generate more stationary time series of fluctuations. We then estimated the pdf of such fluctuations, and looked for an underlying theoretical distribution that fits the results well (Fig. 2). We found a remarkable correspondence between the pdf estimated from the data and the one estimated from the model predictions. The underlying theoretical distribution is strongly non-Gaussian, and consistent with a Lévy alpha-stable distribution with parameter α ≈ 1.3 (Fig. 2). The Lévy alpha-stable distribution is a signature of non-linearities and couplings in complex systems that arise when a system’s components coordinate action and function in unison [35]. We thus see that our model is capable of predicting the statistical properties of measured signals, and that these predictions underpin the hypothesis that beta cells operate collectively.

**Figure 2.**
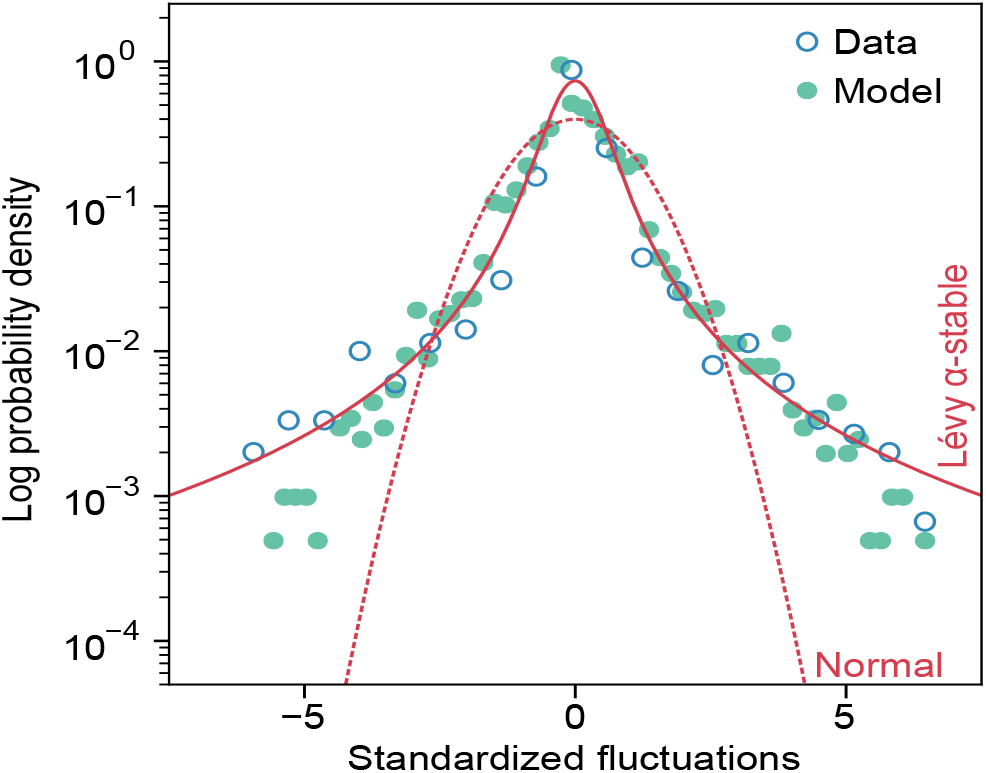
Cell-to-cell communication model predicts statistical properties of measured signals. Denoting the original time series by *X*(*t*), shown are the estimated probability density functions (pdfs) of fluctuations *Z*(*t*) = *X*(*t*) – *X*(*t* – Δ*t*), where Δ *t* = 10 time steps is delay time. Similarity between the pdfs estimated from empirical calcium signals and network model activity is remarkable. Furthermore, the two pdfs are strongly non-Gaussian, which in complex systems such as the studied functional beta-cell syncytium signifies important non-linearities and couplings. The solid curve is a Levy alpha-stable distribution with parameter *α*= 1.3, whereas the dashed curve is a Gaussian distribution with a zero mean and a unit variance.

The link between any node pair in the dynamical network corresponds to the coupling between beta cells. This coupling enables cell-to-cell communication and collective sensing [4], during which a beta cell communicates with its neighboring beta cells through gap junctions and, within a limited range, using paracrine signals [4]. A basic description of the need for communication among heterogeneous beta cells in an islet has been around for some time [36], but has only recently been revived in the light of efficient high-throughput analyses. Such analyses enabled the identification of a number of functional and non-functional cell subpopulations with their characteristic genetic and expression profiles, incidences, and diabetes-related changes [37–39].

Here, we could naturally model the heterogeneity of beta-cell communication channels with the strength of links between neighboring cells. Following the ideas proposed for interneuronal dynamics and synaptic strength [26], in each time step, we strengthened the link between two active nodes in the network by *ϵ* with probability *p_s_*, and weakened by *zϵ* with probability 1 – *p_s_*. If either of the nodes is inactive, we weakened link strength by *zϵ*. Surprisingly, this simple local rule leads to the emergence of a statistical conservation law for the strength of network links as a collective phenomenon [26]. We demonstrated this using the same set of parameters as in Fig. 1E, and setting *ϵ* = 0.001 and *z* = 0.07. We additionally set the initial values of link strengths by drawing randomly from an exponential distribution with unit mean and sufficient standard deviation to capture the heterogeneity of entire cell-cell conductances between pairs of beta cells [40]. When the network is highly active (Fig. 3A), link strengths increase or decrease depending on the activity of individual nodes, creating a wide range of possible outcomes (Fig. 3B). The ensemble average link strength, however, is conserved, thus showing that the collective exhibits a property, namely the statistical conservation of link strength, that is not embedded into any individual node. This modeled property corresponds to the islet-wide conservation of entire cell-cell conductance seen in experiments [21].

**Figure 3.**
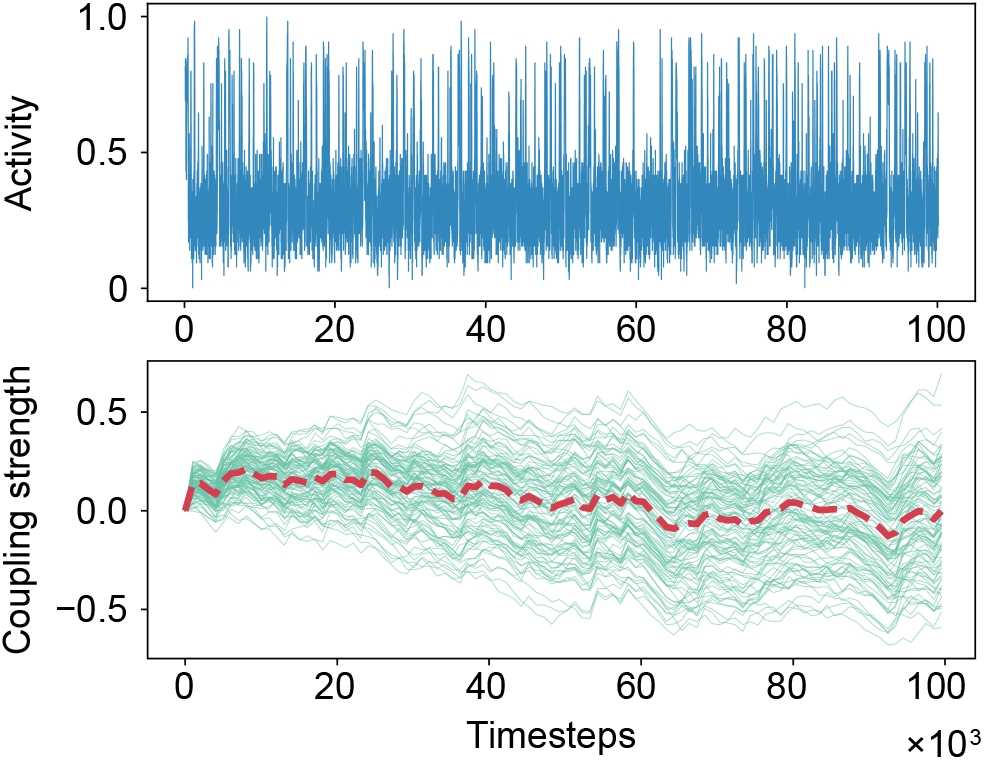
Conserved entire cell-cell conductance of coupled beta cells in stimulatory glucose phase is an emergent collective property. The upper panel focuses on simulated dynamical network activity in the stimulatory phase using the same parameters as in Fig. 1E. In simulations, beta cells are represented by nodes, and cell coupling by links. Each link has a different coupling strength that varies depending on the dynamical network’s activity. The lower panel reveals how the conservation of coupling strength emerges collectively. Coupling between any pair of linked nodes can either decrease or increase in strength (gray solid paths), but the ensemble average (black dashed path) remains unchanged within the limits of statistical fluctuations.

## EMPIRICAL EVIDENCE OF BETA-CELL COLLECTIVE BEHAVIOR

If the response of beta cells to glucose is indeed a collective phenomenon, this should be visible in cell-to-cell communication patterns below and above the threshold glucose concentration. Taking into account that the intensity of cell-to-cell communication is mirrored by the cross-correlation structure of measured signals (Supplementary Remark 2; see also Supplementary Fig. 6), we explored this structure by randomly selecting 4,000 Ca^2+^ traces and calculating cross-correlations

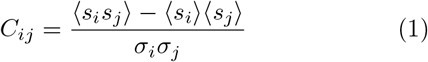

between two traces *s_i_* and *s_j_* at positions *i* = (*x_j_, y_j_*) and *j* = (*x_j_, y_j_*), where *σ_i_* and *σ_j_* are the standard deviations of the traces. To cancel noise, we turned individual cross-correlations into a function of distance *C*(*d*) = (*C_i,j_*), where 〈·〉 indicates averaging over all distances *d_ij_*∈[*d–d*_0_,*d*+*d*_0_] with *d*_0_ = 5 pixels. We calculated *C*(*d*) separately for the non-stimulatory and stimulatory phases, focusing especially on the differences between them.

In the limit of short distances, which are typical distances between the pixels inside a single beta cell, the cross-correlations in both phases are similar. Important quantitative differences emerge at pixel distances above 10. Interpreting function *C*(*d*) as a power-law decay with a faster-than-exponential cutoff

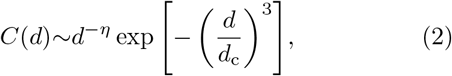

we found that in the stimulatory phase, the correlation scaling exponent is *η* = 0.42 and the characteristic correlation distance in pixels is *d*_c_ = 102 (Fig. 4A). In the non-stimulatory phase, the scaling exponent is lower, *η* = 0.33, and the correlation distance is shorter, *d*_c_ = 90. A shorter correlation distance in the non-stimulatory phase reflects the fact that below the threshold glucose concentration, the collective response of beta cells is still muted.

**Figure 4.**
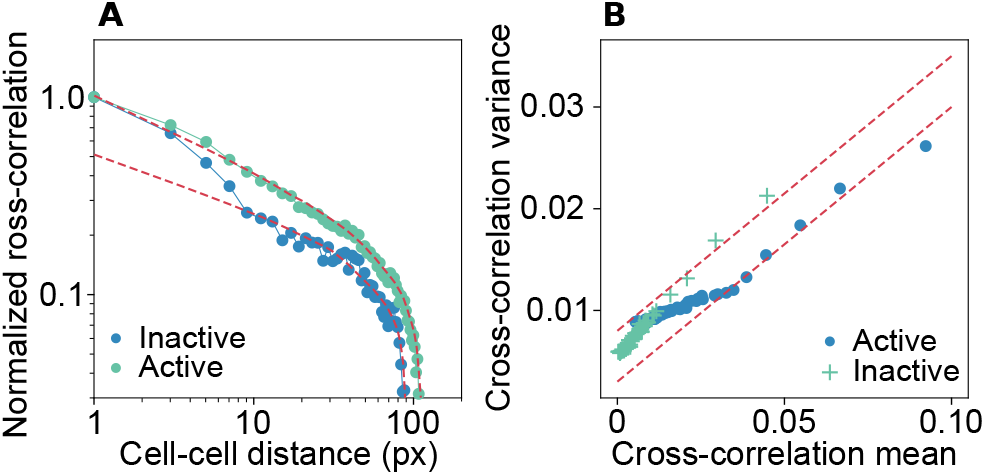
Cross-correlation of Ca^2+^ traces reflects a collective beta-cell response to the threshold glucose concentration. **A**, Shown is ratio *C*(*d*)/*C*(0) in nonstimulatory and stimulatory phases. For a large range of distances, the empirical cross-correlation is interpretable as a combination of power law and faster-than-exponential decay, but with different scaling exponents and correlation distances, mirroring in part a muted collective behavior in the non-stimulatory phase. **B**, Linear relationship between the mean and the variance of empirical cross-correlations is a signature of the Poisson process, yet the process differs below and above the threshold glucose concentration as revealed by two distinct lines that fit the data. This linear relationship starts to break down for smaller cross-correlation means, and thus larger distances at which faster-than-exponential cutoff kicks in (cf. panel A).

Because the cross-correlation function is an expectation of pair cross-correlations, 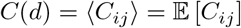, we could also calculate the associated variances, Var [*C_ij_*]. We found that the relationship between Var [*C_ij_*] and 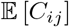 is linear, where two distinct lines corresponding to non-stimulatory and stimulatory phases have the same slope, *c*_0_≈0.25 (Fig. 4B). This result provides additional insights into the nature of fast calcium oscillations. Intuitively, the cross-correlation gets stronger if simultaneous spiking is more frequent and/or larger in magnitude. That Var [*C_ij_*] and 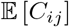 are linearly related is a signature of a Poisson process, thus indicating that the number of simultaneous spikes in two signals recorded at a given distance is a Poission-distributed random variable. We further hypothesize that the number of simultaneous spikes is proportional to the number of gap junctions between nearby cells, in which case slope *c*_0_ is interpretable as an elementary contribution of each cell-cell interaction to the cross-correlation [41], while the estimated value of *c*_0_≈0.25 is in broad agreement with a previous report on collective chemosensing in micropatterned fibroblast cell colonies [41]. The two distinct lines below and above the threshold glucose concentration are yet another sign that cell-to-cell communication within beta-cell collectives qualitatively changes in response to glucose. The distinction between the two phases fades only at large distances where cross-correlations are very weak (bottom-left corner in Fig. 4B).

## DISCUSSION

This work attempts to understand the activity of insulin-secreting beta cells as dynamical networks in which observed oscillatory phenomena emerge from cell-to-cell interactions within the spatial constraints of a typical islet rather than from the properties of single cells alone. In view of a rapid development of electrophysiological probes capable of detecting electrical [42, 43] or optical [44] activities in pancreatic beta cells, a further upgrade of our approach may eventually help to better understand physiology and pathophysiology of beta-cell collectives and yield important cues for diagnosis, therapy, and prevention of diabetes mellitus (Supplementary Remark 3).

A way forward in the context of pathophysiology would be to extract statistical properties of recorded signals, compare them with similar properties of simulated signals, and finally infer the model’s parameter values for which recorded and simulated signals statistically best match each other. Inferred parameter values close to the critical point would then suggest a healthy beta-cell function. Deviations from the critical point, by contrast, would serve as an early warning signal of a potentially degenerative condition (Supplementary Fig. 7).

Particularly in relation to beta-cell transplantation, or better regeneration paradigms in type 1 diabetes mellitus, it is critical to understand to what extent beta-cell collectives have to be replaced or regenerated to reestablish the healthy condition. Much more than just to transplant or regenerate an adequate beta-cell mass, it is important to enable beta-cell collectives to express dynamical cell-to-cell communication ability with a certain relative coupling strength. Our results could thus stimulate the development of novel diagnostic protocols to assess the improving function of beta-cell collectives until full recovery. Although it is too early to decide on the merit of these particular ideas, the future does seem to be in the right intermix of affordable sensing technologies to acquire data, computational-statistical methods to analyze data, and—as much as complexity allows—mechanistic modeling to interpret data.

## Acknowledgements

BP acknowledges financial support from the Slovenian Research Agency (project no. J5-8236), the University of Zagreb’s project Advanced methods and technologies in Data Science and Cooperative Systems (DATACROSS; ref. KK.01.1.1.01.009), and the University of Rijeka. DK, MSK, AS, JD, and MSR received financial support from the Slovenian Research Agency (research core funding program no. P3–0396 and project no. N3-0048). MSR was further supported by the Austrian Science Fund / Fonds zur Förderung der Wissenschaftlichen Forschung (bilateral grants I3562–B27 and I4319–B30). PH was supported by the Japan Society for the Promotion of Science Grant-in-Aid for Scientific Research (B) no. 18H01655. Finally, MJ was partially supported by the University of Rijeka.

## Author contributions

All authors contributed substantially to all aspects of the study.

## Data availability

Raw data analyzed in this study will be made available upon publication through the Open Science Framework repository.

## Footnotes

1 https://github.com/albarji/proxTV

## SUPPLEMENTARY METHODS

### Ethics statement

We conducted tissue isolation for the experiment in strict accordance with all national and European recommendations pertaining to the care and work with experimental animals, including taking all efforts to minimize animal suffering. The protocol was approved by the Veterinary Administration of the Republic of Slovenia (permit number: U34401-12/2015/3).

### Acute pancreas slice preparation

We prepared acute pancreas tissue slices from an 8week old NMRI male mouse following a previously reported procedure [S1, S2]. Briefly, after sacrificing the mouse, the abdominal cavity was accessed via laparotomy. Then, low-melting point 1.9 % agarose (Lonza, USA) dissolved in extracellular solution (ECS, consisting of (in mM) 125 NaCl, 26 NaHCO_3_, 6 glucose, 6 lactic acid, 3 myo-inositol, 2.5 KCl, 2 Na-pyruvate, 2 CaCl_2_, 1.25 NaH_2_PO_4_, 1 MgCl_2_, and 0.5 ascorbic acid) at 38–40°C was injected into the proximal common bile duct, which was clamped distally at the major duodenal papilla. Immediately after the ductal system has been successfully filled, the infused pancreas was cooled with ice-cold ECS and cut out. Tissue slices with a thickness of 140 *μ*m were prepared with a vibratome (VT 1000 S, Leica) and collected in HEPES-buffered saline at RT (HBS, consisting of (in mM) 150 NaCl, 10 HEPES, 6 glucose, 5 KCl, 2 CaCl_2_, 1 MgCl_2_; titrated to pH=7.4 using 1 M NaOH). To allow for staining, slices were incubated for 50 minutes at RT in dye-loading solution (6 *μ*M Oregon Green 488 BAPTA-1 AM (OGB-1, Invitrogen), 0.03 % Pluronic F-127 (w/v), and 0.12 % dimethylsulphoxide (v/v) dissolved in HBS). All chemicals were obtained from Sigma-Aldrich (St. Louis, Missouri, USA) unless otherwise specified.

### Calcium imaging

Individual tissue slices were transferred to a perifusion system containing carbogenated ECS at 37 ^?^C and exposed to single square-pulse-like glucose stimulation with 8 mM (6 mM glucose has been regarded as substimulatory concentration). Imaging was performed on Leica TCS SP5 AOBS Tandem II upright confocal system (20× HCX APO L water immersion objective, NA 1.0). Acquisition frequency was initially set to 10 Hz (512×512 pixels). OGB-1 was excited by argon 488nm laser line and emitted fluorescence was detected by Leica HyD hybrid detector in the range of 500–700 nm (all from Leica Microsystems, Germany) as described previously [S2].

### Data preprocessing

The ensemble-averaged raw signal (Supplementary Fig. 1A) exhibits fast spikes superimposed on slow oscillations, themselves superimposed on a diminishing trend due to dye fading. With focus only on the fast part of the signal, we first used SciPy’s `detrend` function [S3] to remove the gradual amplitude decline (Supplementary Fig. 1B). We thereafter proceeded to separate fast spikes from slow oscillations, to which end we performed signal filtering with proxTV toolbox [S4]. This toolbox is a modern implementation of the total variation denoising technique, which attempts to approximate an original signal with a signal that is “close” to this original (in the sense of a chosen metric), but at the same time minimizes the total variation (and thus noise). Applying this technique returns slow oscillations (Supplementary Fig. 2A), which can then be subtracted from the detrended signal to finally isolate fast spikes (Supplementary Fig. 2B). It is these fast spikes and their statistical properties that we analyzed and modeled in the main text.

**Supplementary Figure 1.**
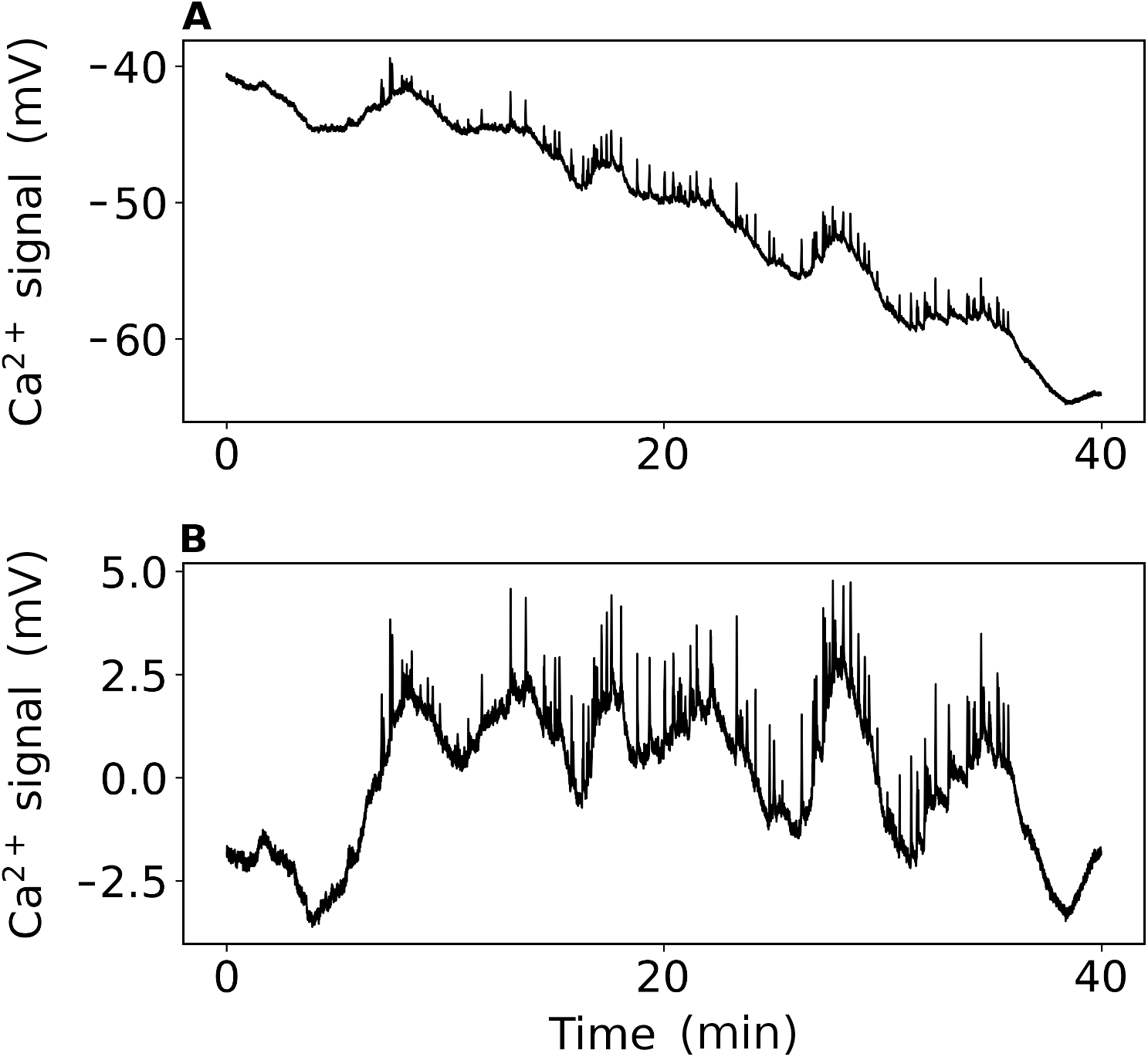
Signal preprocessing I. **A**, Raw ensemble-averaged Ca^2+^ signal showing fast spikes superimposed on slow oscillations, themselves superimposed on a long-term negative trend. **B**, With focus only on fast part of the signal, we first removed the negative trend using SciPy’s `detrend` function.

**Supplementary Figure 2.**
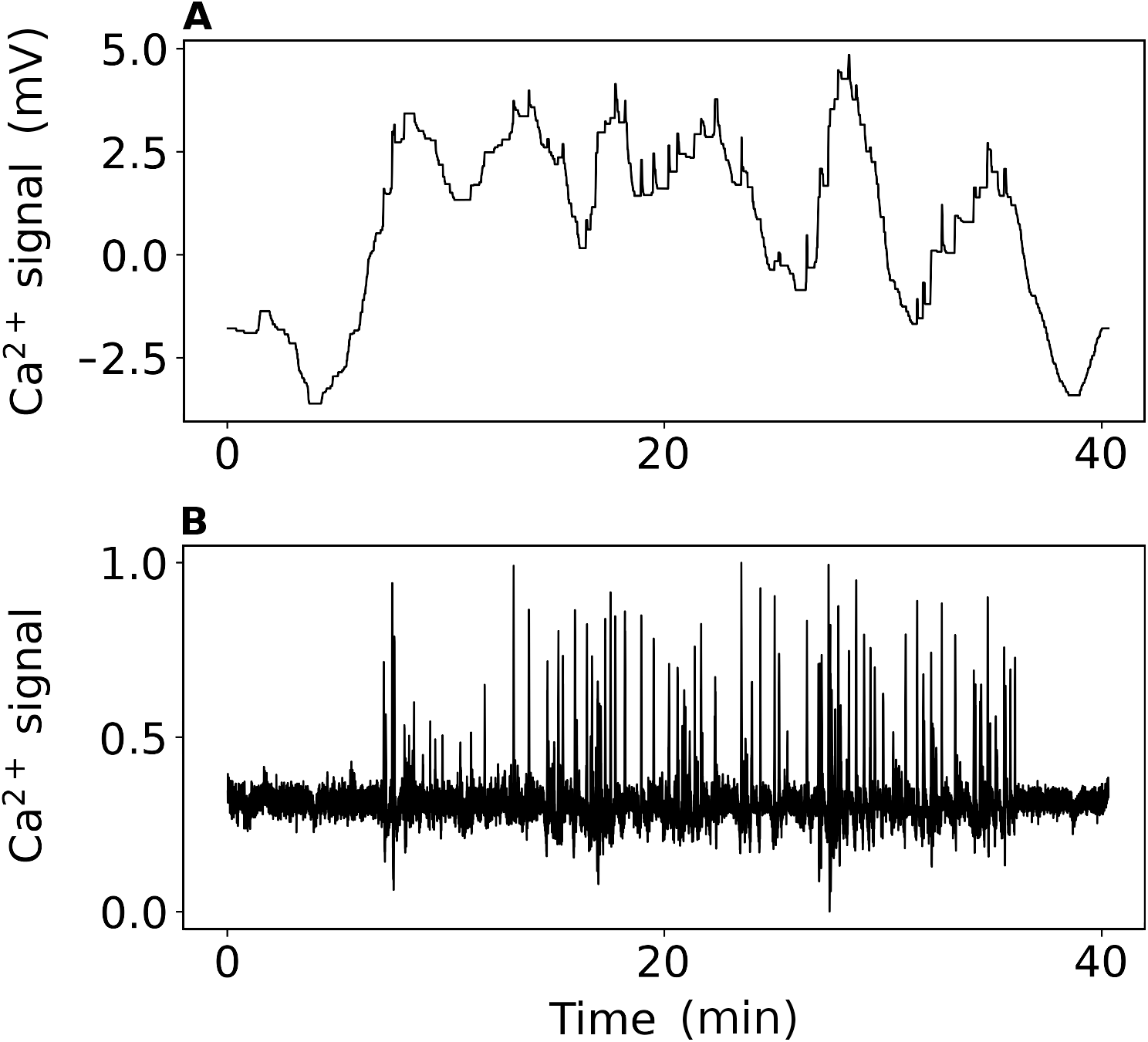
Signal preprocessing II. **A**, Slow oscillations obtained by applying total variation denoising to the detrended signal. **B**, Subtracting slow oscillations from the detrended signal reveals residual fast spikes. The saturation kinetics of the Ca_2_^+^-dependent insulin release suggests that this release is dominated by fast spikes over slow oscillations. It is largely for this reason that we focus on statistical properties and mathematical modeling of the former.

## SUPPLEMENTARY REMARK 1

Here, we present comparisons between the dynamical network model and the empirical data, as well as some additional calculations that characterize the model. The model has five key parameters. These are the node degree, *k*, the threshold number of inactive neighbors, *m*, the inactivation probability rate when *m* or more neighbors are inactive, *r*, the relaxation time after an internal inactivation, *τ*, and the fraction of internally inactive nodes, *p*\* [S5, S6]. For a given network size of *N* = 100 nodes, with node degree *k* = 10 and threshold number *m* = 4, we looked for the values of the remaining three parameters that best correspond to fast Ca^2+^ traces in a statistical sense. Specifically, taking one typical trace (Supplementary Figure 3A) and standardizing it, we find that the skewness is 2.9, while the kurtosis is 12.7. It turns out that by setting *r* = 0.78, *τ* = 10, and *p*\* = 0.28, the dynamical network model produces time series with remarkably similar skewness and kurtosis. These moments for the example in Supplementary Figure 3B are 2.8 and 12.9, respectively.

The dynamical network model is capable of emulating the response of beta cells to glucose stimulation due to node connectivity and mutual, local interactions between node pairs, thus presumably emulating the main characteristics of the beta-cell functional syncytium. More specifically, glucose stimulation of beta cells in experiments corresponds to a decrease in internal node inactivation, *p*\*. For a given external inactivation probability rate, *r*, this implies less suppression from the neighboring nodes. At first the effect is gradual, but near the critical point suppression almost collapses altogether, and network activity picks up instead (Supplementary Fig. 4A). This is in line with empirical evidence that in a coupled configuration, beta cells mute each other’s response to glucose up to a threshold concentration (7 mM in mice), producing a much steeper response curve than in an uncoupled configuration [S7, S8].

From a system dynamics perspective, far from the critical point, the dynamical network model settles in a low-activity equilibrium. As the critical point is approached, however, the high-activity equilibrium becomes more and more influential, which manifests in an increased network activity (Supplementary Fig. 4B). To test the validity of this intuitive explanation, we examined the standard deviation of modeled time series across a range of *p*\* values. Due to a phenomenon known as critical slowing down [S9, S10], we expected to see the evidence of more variation near the system’s critical point. In line with our expectation,the standard deviation indeed spikes close to the critical point, thus confirming that the dynamical network model behaves as we described herein (Supplementary Fig. 5A).

The last property of the dynamical network model examined here is the model’s crosscorrelation structure. We find that the cross-correlation decreases with distance in the network, and generally gets stronger as the model’s critical point is approached from above in terms of the forcing variable, *p*\* (Supplementary Fig. 5B). This is analogous to the measured cross-correlation in beta-cell islets (cf. Fig. 3A in the main text). Namely, the cross-correlation in islets also falls quickly with distance, and generally gets stronger as the threshold glucose concentration of 7 mM in mice is exceeded. As mentioned before, the larger values of glucose concentration correspond to the lower values of the model’s forcing variable *p*\*, thus completing the analogy.

**Supplementary Figure 3.**
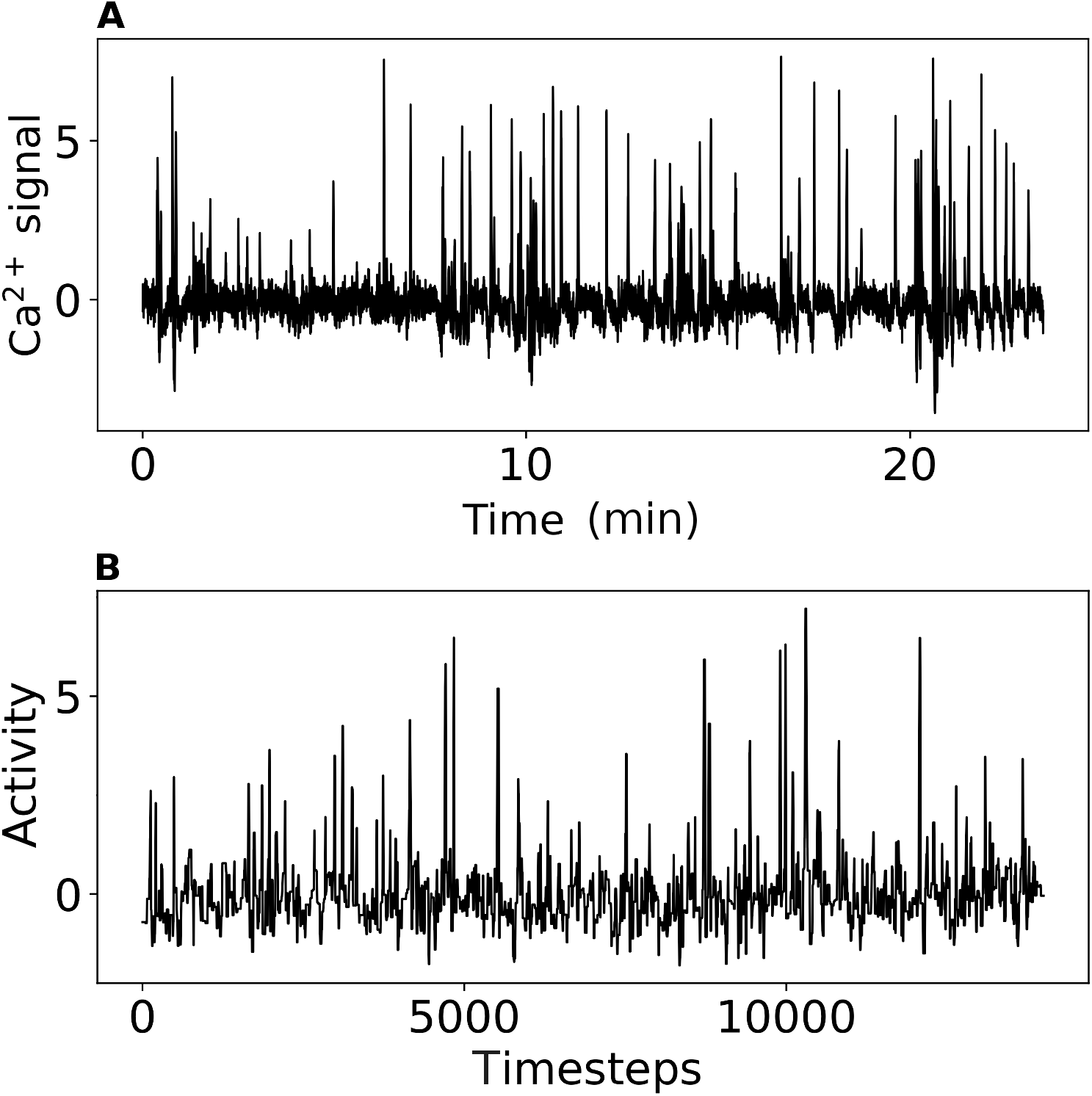
Empirical and model-generated time series have similar statistical properties. **A**, Shown is a typical time series of fast Ca^2+^ signaling after standardization (i.e., the mean is zero and the standard deviation is one). This time series has the skewness of 2.9 and the kurtosis of 12.7. The obtained values are consistent with a standardized time series in which relatively large positive values are frequent. **B**, Here, shown is another standardized time series, but now model-generated with parameters set to *r* = 0.78, *τ* = 10, and *p*\* = 0.28. This time series has skewness and kurtosis very similar to its empirical counterpart above. Specifically, the skewness is 2.8 and the kurtosis is 12.9.

**Supplementary Figure 4.**
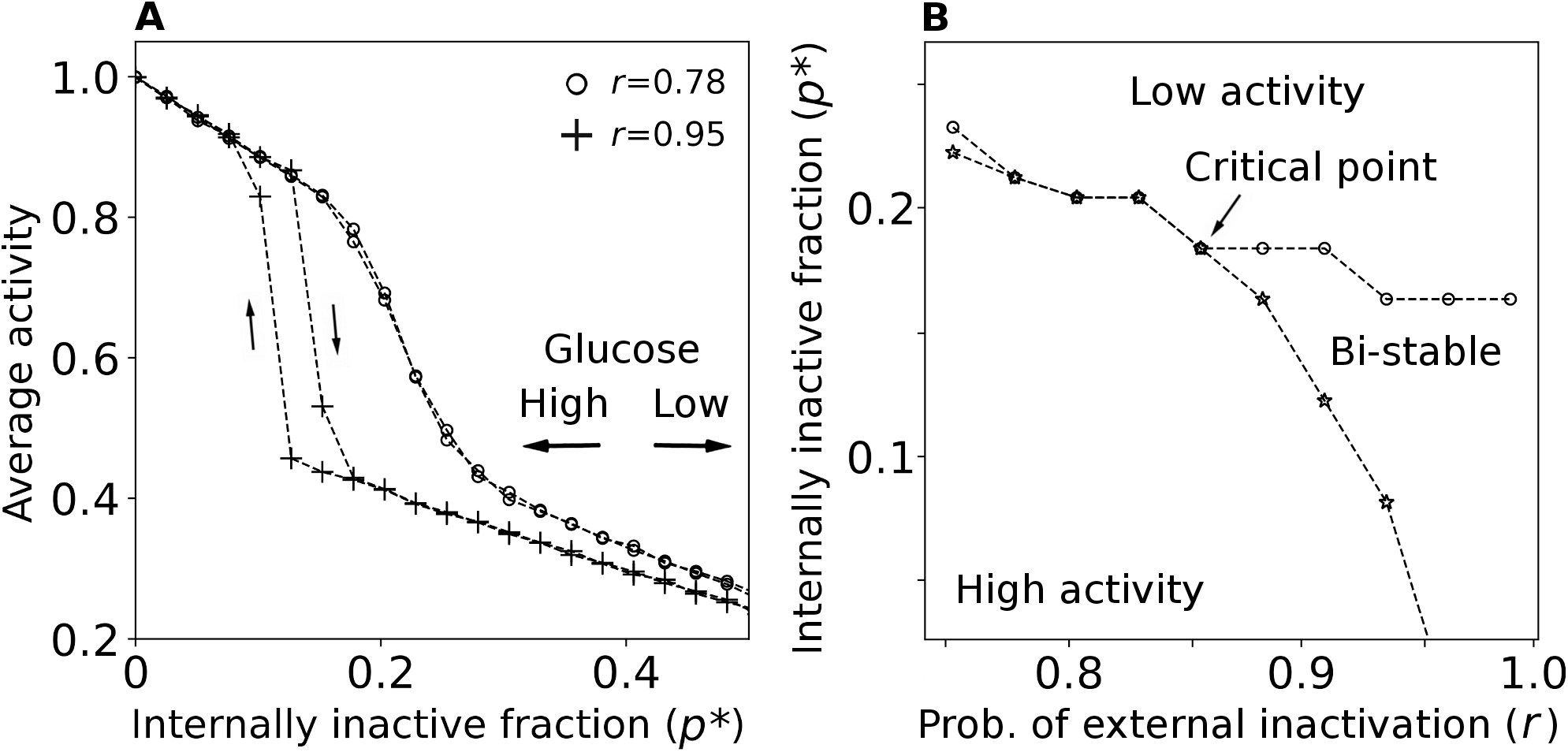
Dynamical network activity increases near the critical point. **A**, Dynamical network model emulates glucose stimulation of beta cells by having the external model forcing, i.e., parameter *p*\*, approach the critical point. Far from the critical point, the dynamical system is under influence of only one, low-activity equilibrium, but near the critical point network activity picks up because of an increasing influence of the second, high-activity equilibrium (see, e.g., [S9, S10]). Shown is the average network activity over many realizations for each value of parameter *p*\* and two fixed values of *r*. When *r* = 0.95, corresponding to very strong cell coupling, a hysteresis loop appears as even spontaneous phase flipping becomes possible. Beta-cell coupling, however, is not that strong because we obtain the best results with values around *r* = 0.78. **B**, Phase diagram of the dynamical network model with three dynamical domains: high activity, low activity, and bi-stable. The curves separating these domains are called spinodals. An increasing glucose concentration in reality corresponds to approaching the (upper) spinodal from above in the model. Near the critical point, in particular, the high-activity equilibrium becomes influential in a sudden, non-linear fashion, generating a more active network in simulations. This effect is a consequence of node connectivity and local mutual interactions between node pairs that mimic beta cells and the beta-cell functional syncytium.

**Supplementary Figure 5.**
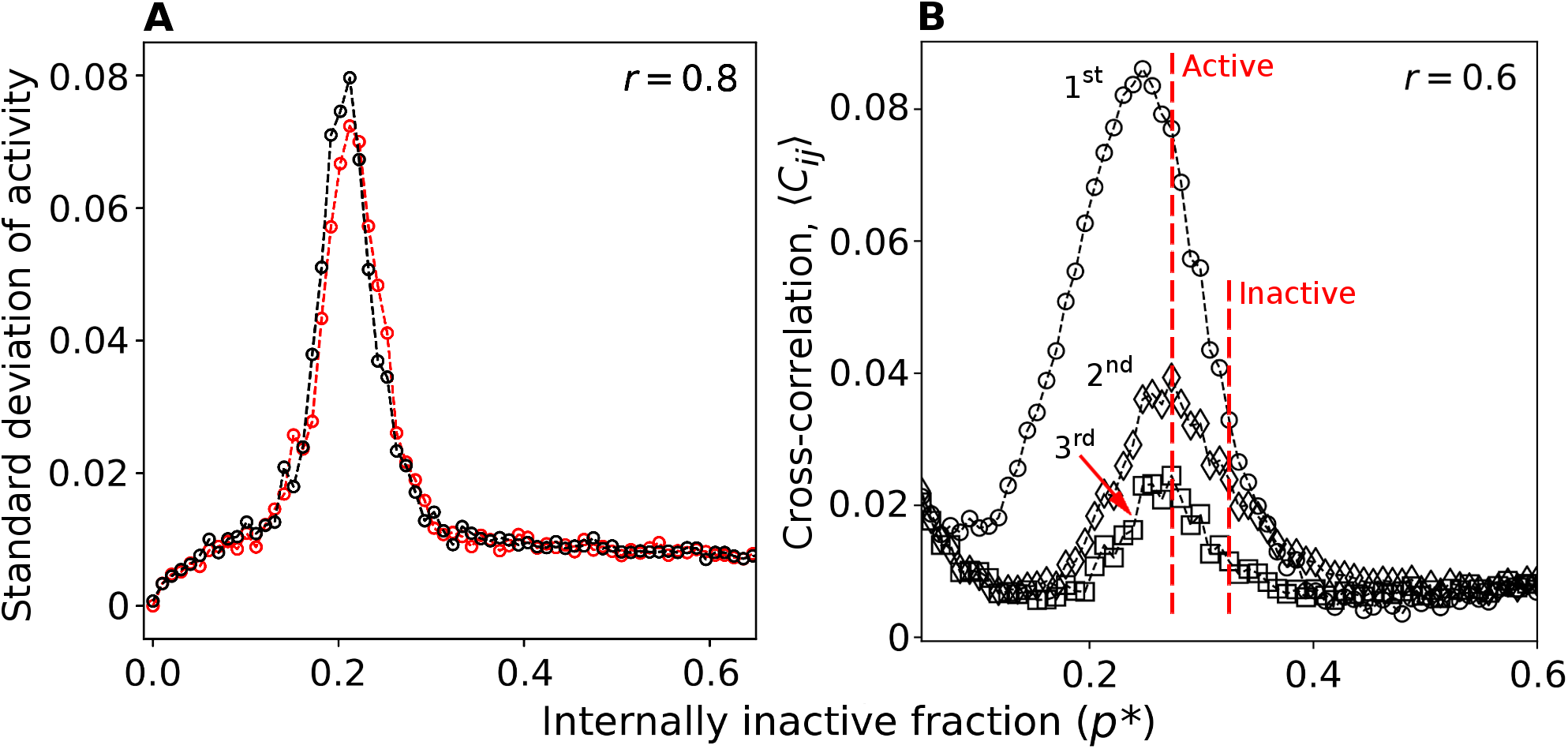
Variation and cross-correlation of network activity change near the critical point. **A**, Shown is the variation of network activity as measured by the standard deviation. The variation spikes near the critical point. This is due to stochasticity that causes the dynamical network model to occasionally flip between the basins of attraction of low- and high-activity equilibria. The black trace corresponds to an increasing glucose concentration (i.e., forcing variable *p*\* decreases), whereas the red trace corresponds to the concentration moving in the opposite direction. **B**, Cross-correlation structure of the dynamical network model is qualitatively similar to the measured cross-correlation in pancreatic islets. Here, shown is the cross-correlation between first, second, and third neighbors in the network. We kept the glucose concentration in the experiments within a relatively narrow range of 6–8 mM, which is just around the threshold concentration for beta cells in mice. In the dynamical network model, however, the upper concentration value of 8 mM should correspond to a value of forcing variable *p*\* close to the model’s critical point, while the lower concentration value of 6 mM should not be too far either (marked by two dashed red vertical lines). The model’s cross-correlation decreases with distance and is generally weaker as the forcing variable increases (i.e., the glucose concentration decreases), which qualitatively corresponds to the measured cross-correlation (cf. Fig. 3A in the main text).

## SUPPLEMENTARY REMARK 2

To determine the extent to which the cross-correlation structure of measured signals mirrors the intensity of cell-to-cell communication we turned to modeling membrane potentials of two coupled beta cells. Pancreatic beta cells are excitable, neuroendocrine cells that undergo periods of quickly oscillating (i.e., spiking) membrane potential followed by periods of silent and slowly changing potential. This dynamics is a subject of intensive studies [S11] that started with Chay-Keizer biophysical model [S12] and a subsequent mathematical analysis by Rinzel [S13, S14].

We reproduced a beta-cell model from Refs. [S15–S17] in Python [S3] following an online example [S18]. At the core of all beta-cell models aiming to generate realistic cell-membrane currents, including the one we implemented here, is the Hodgkin-Huxley model [S19]. The cell membrane is thus described as a capacitor with constant capacitance *C*_m_ and ionic currents flowing through ion channels in the membrane.

Coupling between cells *i* and *j* is due to gap junctions with conductance *g*_c_. Assuming charge neutrality, the equation for the cell membrane potential is

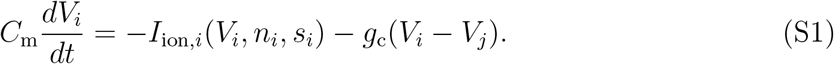

The model incorporates three different ionic currents. Two currents, *I*_Ca_ and *I*_K_, represent the spiking transport of calcium and potassium, respectively. Current *I*_s_ is a slow, inhibitory potassium current that controls bursting behavior. Accordingly

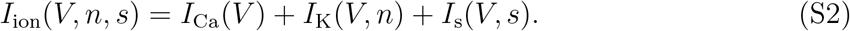

Each ionic current can be written in the form

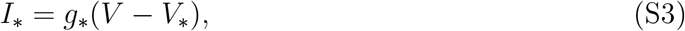

where *g*_*_ is one of the ion-channel conductances and *V*_*_ is the corresponding Nernst potential. More concretely, we have

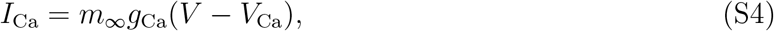

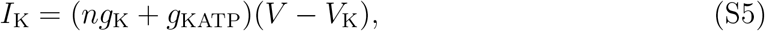

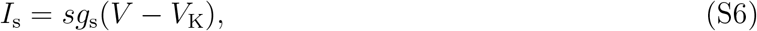

which includes three gating variables, *m, n*, and *s*, to make the ion-channel conductances dynamic. The dynamics of gating variable *m* is, in fact, assumed to be so fast that this variable is always in a voltage-dependent equilibrium, *m*_∞_. The two remaining variables change in time according to

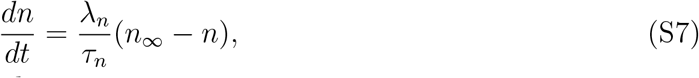

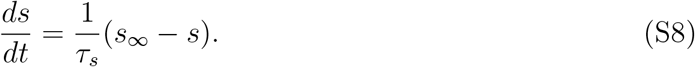

The equilibria of gating variables are given by

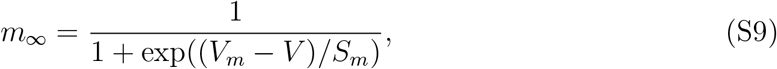

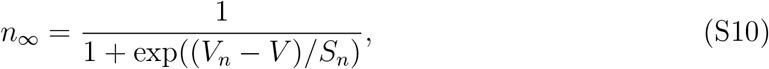

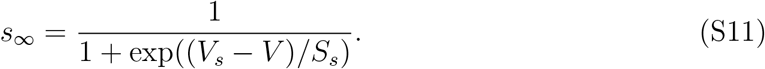

Note that these equilibria attain values between zero and one depending on how cellmembrane potential *V* relates to potentials *V*_*_ and *S*_*_ whose values are defined below. The set of equations can be extended with an equation for intracellular calcium-ion concentration

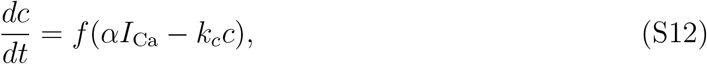

in which case equilibrium value *s*_∞_ and relaxation timescale *τ_s_* become functions of *c* [S20].

To perform computations, we needed to set the parameter values. Starting with betacell membrane capacitance, we used *C*_m_ = 5000 fF and *g_c_* = *g*_KATP_/5, where *g*_KATP_ = 120pS. Parameters for the fast calcium channel were *g*_Ca_ = 1000 pS, *V*_Ca_ = 25.0 mV, *V_m_* = −20.0 mV, and *S_m_* = 12.0 mV. Parameters for the fast potassium channel were *g*_K_ = 2700 pS, *V_K_* = −75.0 mV, λ_*n*_ = 0.95, *τ_n_* = 20.0 ms, *V_n_* = −16.0 mV, and *S_n_* = 5.6mV. Parameters for the slow, inhibitory potassium channel were *g_s_* = 200.0 pS, *τ_s_* = 20000 ms, Vs = −52.0 mV, and *S_s_* = 5.0 mV. The slowness of this channel is reflected in large relaxation timescale *τ_s_* which is three orders larger than the corresponding relaxation timescale *τ_n_* for the fast channel. Finally, parameters for the intracellular calcium-ion concentration were *α* = 4.5×10^-6^ *μ*M fA^-1^ ms^-1^, *k_c_* = 0.2 ms^-1^, and *f* = 0.01. To add high-frequency noise to the cell membrane potential, we perturbed the signal in each timestep with Gaussian noise term e whose mean was zero and standard deviation 0.002.

The described model generates typical bursting oscillations in the form of a steady alteration between spiking and silent states (Supplementary Fig. 6A). For the present study, it is crucial that the cross-correlation coefficient between the two shown membrane potential traces depends on the gap junctional conductance. Specifically, after setting up the model’s parameters such that weakly coupled cells exhibited bursting (Supplementary Fig. 6B), we started increasing the relative coupling strength (i.e., the ratio of gap junctional to K_ATP_-channel conductances), and recording said cross-correlation coefficient. For weakly coupled cells, we found that the cross-correlation increases linearly with coupling strength until the coupling turns strong, and cells become fully synchronized (Supplementary Fig. 6C). The obtained linear relationship indicates that more strongly coupled cells (i.e., those having a better means of cell-to-cell communication) also produce more strongly correlated signals. An immediate implication is that the estimated cross-correlation is a reflection of cell-to-cell communication.

**Supplementary Figure 6.**
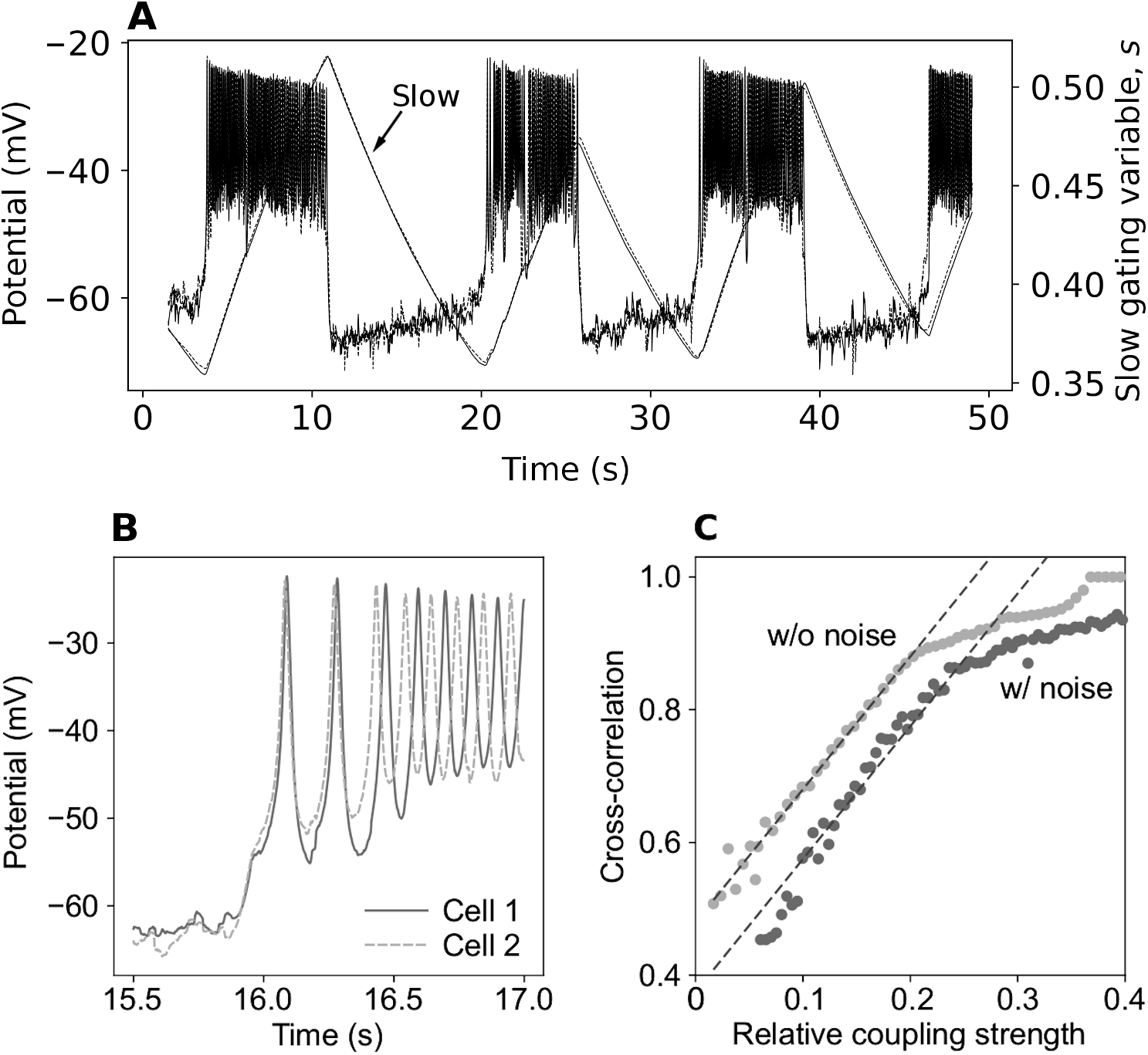
Cross-correlation between electric potentials of coupled beta cells is dependent on the gap junctional conductance. We obtained this result using a well-known mathematical model of beta-cell electrical activity described in the text. **A**, Shown are the membrane potential and the level of the slow gating variable as the functions of time. Model successfully generates typical bursting oscillations in the form of a steady alteration between spiking and silent states. **B**, Zooming into the plotted signals reveals that the model accounts for the electrical activity of two coupled beta cells. A sure tell sign of weak coupling is the phase shift that develops between the potentials during the burst. **C**, Cross-correlation of the membrane potentials of two weakly coupled beta cells, computed with and without Gaussian noise current. The cross-correlation coefficient is indeed proportional to the gap junctional conductance (relative to K_ATP_-channel conductance) until saturation at large coupling strengths.

## SUPPLEMENTARY REMARK 3

Dynamical networks offer means to study damage accumulation and the consequences of such damage on network operation [S21]. In the context of the present study, damage accumulation by which individual nodes become unable to enter, e.g., the active state, would correspond to the progression of a disease similar in nature to type 1 diabetes mellitus. To demonstrate the effects of damage accumulation on network operation, particularly on the collective mode that is presumably vital for glucose homeostasis, we computed the average network activity after a period of transient dynamics during which nodes that turn inactive may stay locked in that state. Specifically, when a node is in the inactive state, the relaxation time of this node may change with probability q from the default value of *τ* = 10 to *τ* ≫ 10. The node thus stays permanently inactive throughout a finite lifetime simulation.

The results of permanent node inactivation are striking. At first, the dynamical network responds to damage accumulation with a linear decrease of activity in time (Supplementary Fig. 7A). When the critical amount of damage accumulates, however, the non-linear loss of activity is sudden and large, i.e., the network’s collective mode of operation becomes compromised (Supplementary Fig. 7A). Furthermore, the average network activity may severely diminish through the course of a finite lifetime, provided that the value of parameter q is large enough (Supplementary Fig. 7B). Translating these results into a better understanding of the collective beta-cell operation, we see that a lowered average network activity may correspond to the onset of a disease such as type 1 diabetes. This disease is indeed known to arise from beta-cell dysfunction such that postprandial increases in blood glucose concentration can no longer be counteracted. Armed with these concepts, we conclude that devising cheap, effective, and unobtrusive diagnostic techniques is a matter of complementing reasonable technological advances with well-established statistical methods. The technology in question is high-fidelity data loggers to monitor and record beta-cell activity. Signals thus recorded could then be subjected to statistical analyses to detect the early signs of trouble, recognizable as deviations from normal beta-cell functioning in the collective mode.

**Supplementary Figure 7.**
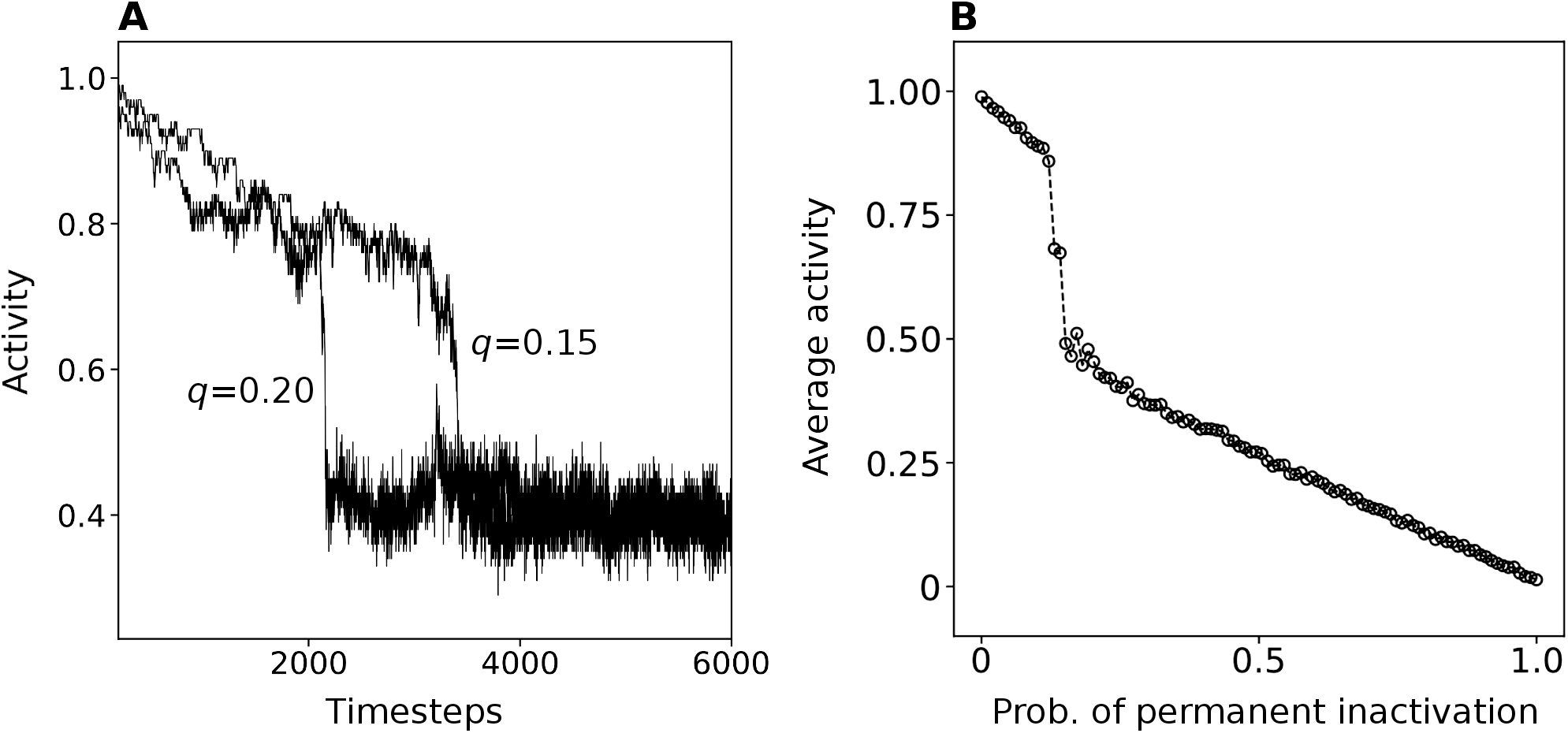
Damage accumulation leads to a failure of the collective network mode. Shown is a stylized situation in which the dynamical network is highly active (*p*\* = 0.01, *r* = 0.90, and *τ* = 10), but whenever a node turns inactive, there is a good probability, denoted *q*, that the inactivation is permanent. Such inactivation would correspond to beta-cell dysfunction reminiscent of the progression of type 1 diabetes. **A**, Damage accumulation is slow at first, leading to a linear decrease of the dynamical network’s activity over time. This changes as the critical amount of damage is accumulated, in which case the network suddenly loses the ability to activate a large proportion of nodes, i.e., the network’s collective mode of operation is compromised. Technically, the damage keeps accumulating thereafter as well, but at a very slow pace. **B**, Here, we show the average network activity after a period of transient dynamics, i.e., calculated using timesteps 5000–6000, as a function of the permanent inactivation probability, *q*. When this probability is small, the dynamical network remains largely operational over long periods of time. Increasing q values correspond to faster damage accumulation, which is why, for large enough *q*, the failure of the collective mode becomes inevitable within the given time window. This is evidenced by a steep drop in the average network activity around *q* = 0.12.

